# Salinity shock in *Jatropha curcas* leaves is more pronounced during recovery than during stress time

**DOI:** 10.1101/378208

**Authors:** Leonardo Silva-Santos, Natália Corte-Real, Jaqueline Dias-Pereira, Regina C.B.Q. Figueiredo, Lauricio Endres, Marcelo F. Pompelli

## Abstract

To verify the possible morphological and ultrastructural differences in the *Jatropha curcas* leaves, in response to high-intensity salt stress, three genotypes were evaluated (CNPAE183, JCAL171 and CNPAE218). In all the genotypes, 750mM NaCl, added to the nutrient solution, was applied to test its salt tolerance. For the analysis, the leaves were collected at three time points: (*i*) before stress (time 0 hour); (*ii*) during stress time (time 50 hours); and (*iii*) in the recovery period (time 914 hours) when the stressed plants recovered and demonstrated measurements of net photosynthetic with values similar to those demonstrated by the control plants. We showed that regardless of the genotype, saline shock caused an increase in the thickness of the mesophyll, and after the removal of NaCl, the thicker mesophyll remained in the JCAL171 and CNPAE218 genotypes, while the values observed in the CNPAE183 genotype were similar to those before stress. Scanning electron microscopy indicated that the stomata of CNPAE183 are smaller and have a stomatal index higher than the values demonstrated in JCAL171 and CNPAE218. Therefore, among the genotypes analysed, CNPAE183 demonstrates that it could be considered a promising genotype for future studies of genetic improvement that seek elite genotypes tolerant to salinity.

**Highlights:** This manuscript present the following highlights:

The mesophyll thickness contributes to provide a smaller path for the CO2 to Rubisco *J. curcas* may reduce mesophyll air spaces as a strategy to mitigate low gas exchange Leaves modulate the expansion of stomata differently than other epidermal cells Smaller stomata with greater pore aperture are more abundant on the abaxial surface CNPAE183 is a candidate for studies in search of elite genotypes tolerant to salinity

## Introduction

The physic nut (*Jatropha curcas* L.) is a woody-shrub species belonging to the Euphorbiaceae family. Its true centre of origin is unknown, but studies indicate that the species is native to Central and South America [1]. In recent years, this species has been researched for exploitation of the commercial biodiesel production [2, 3]. *J. curcas* is easily propagated and has a short period of growth until the first fruit harvest [4], low seed cost [2], high oil content (40–58%) [5, 6], and good adaptation to different agroclimatic conditions [7–9], thus contributing to the wide geographical distribution of the plant, which is present in practically all the intertropical regions of the world [9, 10]. The physiological characteristics of the species, associated with its economic potential, can transform *J. curcas* into an alternative biofuel plant for arid and semiarid regions [11]. However, the scarcity of rainfall and the high evaporative demand of these regions induce a high concentration of salts in the soils, a situation that is exacerbated by the use of brackish water for irrigation [12].

It is known that the high concentration of salts in the soil solution represent a stress factor for plants [13] related to the action of the ions in the cytosol and the reduction of the osmotic potential [14], causing osmotically retained of water in the saline solution, decreasing the water availability for plants [15]. Salinity stress (*i.e*., salinity shock or salinity for a moderate to long time) causes various effects on the plant, such as (*i*) limiting plant growth [16–18], (*ii*) a stomatal closure [15], (*iii*) a reduction in root and stomata conductivity [19], (*iV*) an increase in succulence and leaf thickness [18, 19], (*v*) a foliar injury and abscission [20] and (*vi*) a reduction in net photosynthesis [17, 18]; factors that strongly reduce the productivity of the cultivated plants and have a greater effect than any other phenomenon [14, 21]. Salinity shock is usually more intense and leads to abrupt changes in plant physiology, causing phenotypic changes that may or may not be apparent. Phytoene synthase (*psy*) and phytoene desaturase (*pds*), genes closely related to the carotenoid synthesis, are reported to decrease under salinity up-shock, leading to a decrease in carotenoid synthesis or chlorosis [22] in the leaves. It is very likely that if the chlorophyll synthesis is affected, all the pathways that depend on it are also decreased. These include gas exchange, growth and development and foliar expansion, and physiological and morphological features. Salinity shock controls their overall metabolic activity and directs it towards adaptation to the imposed stress. The adapted plants could manage stress totally or partially, while non-adapted plants will escape stress or die. The adaptability of the species studied to the effects of salinity shock has resulted in low variation in their water content, which dependes primarily on their ability for internal osmoregulation [15].

The performance of *J. curcas* under salt stress has been the target of several physiological studies. However anatomy studies of this species have rarely been performed. Many researchers consider *J. curcas* to be a species sensitive to salinity [14, 15], while others classify it as moderately tolerant to salt stress [13]. Another researchers considers *J. curcas* to be a halophyte-like plant [23]. Such contrasting results can be explained by the fact that *J. curcas* is still in the domestication stage [24]. Several studies that examined the genetic variability of the species have been developed in the last few years [1, 3, 25], providing promising results that interact to generate more productive cultivars that are more closely adapted to the regional edaphoclimatic conditions, thus consolidating the species as an alternative for biodiesel production [10, 13]. This study aimed to study the morphological, anatomical and ultrastructural features affected by the salinity shock in three *J. curcas* genotypes, how the genotypes perceives salinity shock and how recovery occurs after the alleviation of saline conditions, mimicking what can happen in nature in commercial plantations irrigated with brackish water.

## Material and Methods

### Study site, plant material and salt stress input

The experiment was carried out in a greenhouse at Federal University of Pernambuco (8°02′60”S, 34°56′52”W, 18 m a.s.l). After previous studies of germination, growth and gas exchange under salinity, three genotypes from different regions of Brazil (Table 1) with contrasting responses to salinity were selected: CNPAE183, JCAL171 and CNPAE218; these genotypes are tolerant, moderately tolerant and sensitive to salinity, respectively (Corte-Real, N.; manuscript in preparation). Seeds of these three genotypes were germinated until the cotyledons opened in trays with a 5 L capacity that containing washed river sand. After that, the seedlings were individually transplanted into 9 L pots containing washed river sand and irrigated every two days with 50% Hoagland solution [26], where they remained for 15 days for acclimatization. Subsequently, the nutrient solution was replaced by 100% Hoagland solution, and the plants remained in this condition for three months when they reached a suitable height for the start of the experiment (time 0 hour). From this moment, the salinity shock treatment began. The seedlings were watered at approximately 7:00 am, with 800 mL of 100% Hoagland solution plus 750 mM L^−1^ of NaCl, corresponding to 47 dS m^−1^ of electrical conductivity. After the stress-promotion time (*i.e*., 50 hours), the soil was thoroughly washed with distilled water to remove the salinity from the soil solution (*i.e*., soil with electrical conductivity ≤ 3 dS m^−1^). At this time, the plants received 100% Hoagland solution for an additional 35 days (914 hours), when the gas exchange parameters of the stressed plants reached the same level as the control plants (Corte-Real, N.; manuscript in preparation).

**Table 1.** Information and location of the three genotypes of *Jatropha curcas* studied under salinity conditions.

| Genotype | City | State | Location | Altitude (m.asl) |
| --- | --- | --- | --- | --- |
| 183 | Jaíba | Minas Gerais | 23°47'55.0" S 53°18'48" W | 478 |
| 171 | Maceió | Alagoas | 09°27'60.0" S 35°49'41" W | 131 |
| 218 | São Miguel do Araguaia | Goiás | 13°55'57.0" S 50°09'17" O | 350 |

#### Optical and ultrastructural leaf anatomy

Leaf fragments from the interveinal regions of fully expanded healthy leaves were collected from the middle third of the plants for the anatomical evaluations. The leaf fragments were collected at three different times: (*i*) before stress (*i.e*., 0 hour), (*ii*) stress-promoting time (*i.e*., 50 hours), and (*iii*) at recovery (*i.e*., 914 hours). At each respective time, the leaf samples were collected, readily fixed in FAA50% [27] and transferred to 70% alcohol until use. The samples were then dehydrated in graded ethylic series, included in plastic resin (Leica Historesin Embedding Kit, Leica Microsystems, Wetzlar, Hesse, Germany), cross-sectioned (7 μm) in an automated microtome (Microtome, mod. RM 2255, Leica Microsystems Inc., Deerfield, IL, Wetzlar, Hesse, Germany) stained with Toluidine Blue [28] and permanently mounted on synthetic resin (Permount^™^ Mounting Medium, Fisher Chemicals, Sunnyvale, CA, USA). All the materials were photographed using a microscope (model DM750, Leica Microsystems, Wetzlar, Hesse, Germany) coupled to an HD camera (model ICC50 HD Leica Microsystems, Wetzlar, Hesse, Germany). From these images, the thickness of the periclinal wall of both epidermis surfaces and palisade and spongy parenchyma, as well as, the area of air spaces in the mesophyll were measured using Image-Pro Plus software (Version 4.5.0.29, Media Cybernetics, Silver Spring, MD, USA).

Scanning electron microscopy (SEM) was used to count the stomata and study the stomatal structure in detail. For this, 5 mm fragments of the leaves were fixed in Karnovsky buffer [29] for 30 days. The fragments were washed three times with cacodylate buffer (sodium cacodylate trihydrate, part number C0250, Sigma Aldrich, St. Louis, USA), dehydrated in a graded ethyl series, dried to a critical point (Bal-Tec CPD 030 critical point dryer, Bal-Tec, Balzers, Liechtenstein, Germany), covered with a thin gold (Metalizer, Denton Desk II Sputter Coater, Torontech Group International, Markham, ON, Canada) and photographed under a scanning electron microscope (Scanning Electron Microscope, mod. JSM-5600LV, Jeol, Peabody, MA, USA). To measure the stomatal density and stomatal index, we used the Salisbury equations [30] with modifications proposed by Pompelli, Martins [31]. Measurements of the stomata length, width and area were performed as described by Pompelli, Martins [31], while the stomatal pore aperture was performed by direct measurement of the images.

#### Leaf area and specific leaf area measurements

For the leaf area measurement, 20 healthy and fully expanded leaves were collected from four plants cultivated without the addition of salt. The leaves were then scanned using a scanner (mod. HP Scanjet G2410 1,200 x 1,200 dpi, Hewlett-Packard Corporation, Palo Alto, Ca, USA) and measured using Image-Pro Plus software (version 4.5.0.29, Media Cybernetics, Silver Spring, MD, USA). In each leaves, 40 discs (~1.5 cm^2^) were collected from random points of the leaf surface in order to determine the specific leaf area (SLA), which was calculated by the ratio between the leaf area and the leaf mass [32].

#### Statistical analysis

The experimental design was completely randomized, with four replications and three accessions. The results were submitted to a two-factor ANOVA and a Student-Newman-Keuls test (α = 5%). The analyses were performed using R programming language [33].

### Results

#### Morphoanatomical characterization of the J. curcas genotypes

The three genotypes evaluated (CNPAE183, JCAL171 and CNPAE218) demonstrated petiolate and alternate leaves with simple or palm-shaped leaf morphology, containing 5 to 7 lobes each (Supplementary Fig. 1). The architecture of the non-stressed plants demonstrated a similar pattern, independent of the genotype evaluated; however, the development of the plants grown under salinity shock was highly reduced, with the genotype CNPAE218 being the most affected by salinity (Supplementary Fig. 1). These findings became even more pronounced after stress, since at this time, the plants that were under salinity, aborted a significant portion of their leaves, although this was reversed with alleviation of the saline conditions. CNPAE183 demonstrated as the fastest plant recovery, with the regrowth of new leaves in a few days. All the plants submitted to salinity strongly reduced their final height by ~20% when compared to the non-stressed plants, regardless of genotype. Genotype CNPAE183 showed the highest unitary leaf area among the evaluated genotypes, *i.e*., 46% higher than the JCAL171 genotype and 64% higher than the CNPAE218 genotype (Fig. 1). The specific leaf area (SLA) did not differ significantly between the genotypes and neither in response to salinity shock.

**Figure 1:**
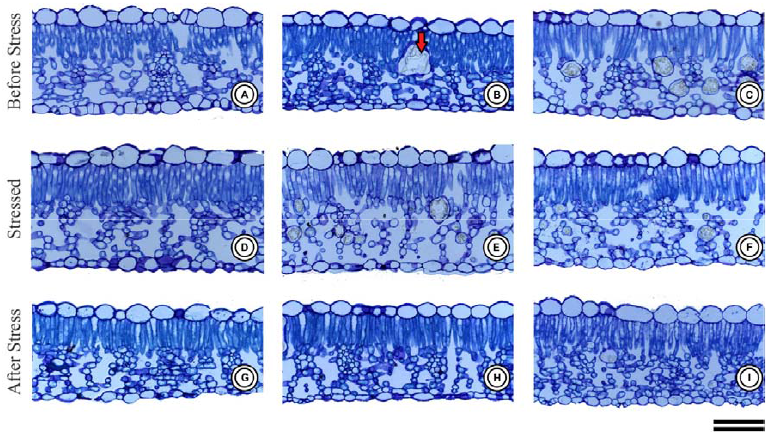
Photomicrograph of cross-sections of leaves of the three different genotypes of *Jatropha curcas* (CNPAE183, A, D and G; JCAL171, B, E and H and CNPAE218, C, F and I), subjected to 750 mM NaCl for 50 hours (D-F) and recovery after 35 days after salt stress alleviation (G-I). Bars = 100 μm. Red arrow denote drusen, which repeat many times in the spongy parenchyma.

#### Morphological characterization of the leaves under saline conditions

The morphoanatomy of *J. curcas* revealed the presence of a uniestratified epidermis, with smooth and thin-walled cells with rounded contours and irregular shapes. The adaxial surface of the epidermis was ~59% thicker than the abaxial surface (Fig. 1 and Table 2) on which a thin layer of wax is deposited. The thickness of the epidermis (adaxial and abaxial surfaces) of genotype CNPAE183 was not affected by salinity shock. Alternatively, the epidermis thickness, on both surfaces, of the JCAL171 and CNPAE218 genotypes was reduced in the leaves after stress. In these genotypes, the epidermal thickness of the adaxial surface decreased by 13.6% and 16.3%, respectively, while the reduction of the epidermal thickness of the abaxial surface was decreased by ~16% in these two genotypes (Table 2).

**Table 2.** Anatomical traits [upper and lower epidermis thicknesses, palisade and spongy parenchyma thicknesses, air spaces in palisade (air PP) and spongy (air SP)], in the leaves of *Jatropha curcas* subjected to 750 mM NaCl in three events (before stress, stressed and after stress). Different upper-case letters represent statistical significance between the means for genotype and time, and different lower-case letters represent statistical significance among the genotypes in the different timeline (*P* ≤ 0.05, Newman–Keuls test). Asterisks denote statistical significance among the means in the adaxial and abaxial surface in the epidermal thickness, genotype and timeline (*P* ≤ 0.001, t-test). The values represent the media (± SE). n = 4

| Timeline | Epidermal thickness ($\mu\text{m}$) | | Mesophyll thickness ($\mu\text{m}$) | | Air Space (%) | |
| --- | --- | --- | --- | --- | --- | --- |
|  | Adaxial | Abaxial | Palisade tissue | Spongy tissue | Air PP | Air SP |
| Before stress (time 0 hour) |  |  |  |  |  |  |
| CNPAE183 | 34.7 ± 1.1 Aa* | 21.4 ± 0.7 ABa | 60.4 ± 0.9 Ab | 90.8 ± 2.6 Ab | 7.6 ± 0.7 Aa | 33.5 ± 1.4 Aa |
| JCAL171 | 33.5 ± 1.0 Aa* | 22.5 ± 0.5 Aa | 58.2 ± 1.0 Ac | 85.0 ± 1.5 Ab | 6.3 ± 0.6 Aa | 32.8 ± 1.7 Aa |
| CNPAE218 | 33.6 ± 1.1 Aa* | 20.0 ± 0.9 Ba | 60.4 ± 1.0 Ab | 89.6 ± 1.9 Ab | 6.9 ± 0.8 Aa | 32.0 ± 1.8 ABa |
| Stressed (time 50 hours after start stress) |  |  |  |  |  |  |
| CNPAE183 | 34.0 ± 0.7 Aa* | 20.5 ± 0.8 Aa | 66.4 ± 1.4 Aa | 103.6 ± 3.1 Aa | 7.8 ± 0.72 Aa | 34.9 ± 1.7 Aa |
| JCAL171 | 31.1 ± 0.8 Aab* | 21.3 ± 0.7 Aa | 66.9 ± 1.2 Ab | 93.5 ± 4.1 Ab | 4.6 ± 0.5 Bb | 26.7 ± 1.2 Bb |
| CNPAE218 | 34.4 ± 1.0 Aa* | 22.1 ± 0.4 Aa | 64.8 ± 0.8 Aa | 100.7 ± 3.8 Aa | 4.3 ± 0.5 Bb | 23.2 ± 1.0 Bb |
| After stress (Recovery, 914 hours after start stress) |  |  |  |  |  |  |
| CNPAE183 | 32.8 ± 0.3 Aa* | 20.4 ± 0.5 Aa | 61.3 ± 1.0 Cb | 87.3 ± 2.1 Bb | 6.6 ± 0.7 Ab | 31.7 ± 0.7 Ab |
| JCAL171 | 29.5 ± 1.1 Bb* | 19.4 ± 0.6 Ab | 72.0 ± 0.8 Aa | 111.0 ± 4.7 Aa | 5.2 ± 0.3 Bb | 31.0 ± 1.3 ABa |
| CNPAE218 | 28.9 ± 0.4 Bb* | 17.2 ± 0.6 Bb | 65.3 ± 1.57 Ba | 103.8 ± 2.7 Aa | 4.3 ± 0.42 Bb | 27.9 ± 0.7 Bb |

*J. curcas* leaves demonstrate a dorsiventral mesophyll arrangement, with a uniestratified palisade parenchyma with elongated and compacted cells that face the adaxial surface and a portion of spongy parenchyma, facing the abaxial surface, consisting of a superposition of 7 to 10 layers of cells of irregular and small compacted sizes, resulting in many intercellular spaces with a large amount of drusen (Fig. 1, red arrow). Salinity shock caused a strong increase in the thickness of the palisade (28.8%) and spongy (30.5%) mesophyll in the leaves of the JCAL171 genotype; a higher increase was observed in the leaves following the alleviation of the saline conditions, while this increase was slightly lower in the genotype CNPAE218 (*i.e*., 8.1% and 15.9%, for palisade and spongy mesophyll, respectively). Unlike the other genotypes, CNPAE183 tended to increase the thickness of the mesophyll during stress, but this condition did not translate into the new leaves that emerged after the alleviation of the saline condition (Table 2).

The air spaces in the mesophyll were genotype- and treatment-dependent, since there was interaction between these factors. The genotype CNPAE183 had fewer air spaces in the mesophyll, demonstrating a reduction of the air spaces in the palisade and spongy parenchyma of the 13% and 5%, respectively. JCAL171 demonstrated a reduction of the air spaces in the palisade and spongy parenchyma of 38% and of 23.1%, respectively, and genotype CNPAE218 demonstrated a decrease of 38% and 13% in the air spaces in the palisade and spongy parenchyma, respectively.

#### Stomatal features

*J. curcas* demonstrated amphistomatic leaves with paracytic-like stomata abundantly distributed throughout the epidermis (Fig. 2). Of all of the genotypes evaluated, the abaxial surface has a stomatal density 6 times higher than observed in the adaxial surfaces (Table 3, Fig. 2). In addition, the stomata recorded on the abaxial surface are abundantly uniformly distributed along the surface, while on the adaxial surface they are sparse and concentrated in the regions close to the ribs. Genotype CNPAE183 was the only genotype evaluated that showed a significant increase in the stomatal density in the leaves after salinity shock, both in the adaxial (65%) and in the abaxial surfaces (23.4%). Even though, the stomatal density of the abaxial surface was 6 times higher than that observed in the adaxial surface, the stomatal area did not exhibit the same tendency. While the stomatal area of the abaxial surface was ~9% smaller than the stomatal area of the adaxial surface in the genotype CNPAE183, in the genotypes JCAL171 and CNPAE218, the stomata area of the abaxial surface was ~9.1% and 8.5% higher in the abaxial surface, respectively (Table 4). However, this decrease or increase in the stomatal area did not translate into significant differences in the linear dimensions of the stomata (*i.e*., both the stomatal length and the stomatal width did not change under salinity shock), except for the JCAL171 genotype that demonstrated a stomatal width 7% greater than those that emerged after the alleviation of the saline conditions (Table 3). Regardless of the treatment, the stomatal aperture of the abaxial surface was 6 times higher than that observed in the adaxial surfaces (Table 3, Fig. 2). Despite this, the adaxial stomatal aperture was not significantly affected by salinity shock treatments in any genotypes evaluated. In distinct form, the abaxial stomatal aperture was significantly decreased in CNPAE183 (6%) and JCAL171 (4%), while no significant difference (P = 0.387) was perceived in CNPAE218 in the leaves that emerged after the alleviation of the saline conditions (Table 4).

**Figure 2:**
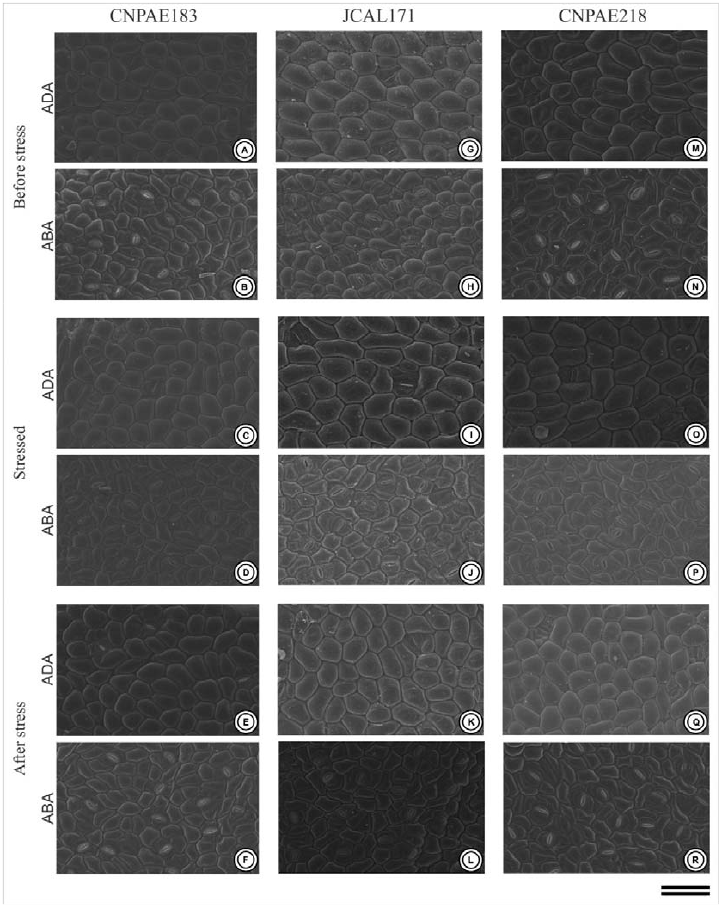
Electron scanning microscopy of the leaf surface of the three different genotypes of *Jatropha curcas* (CNPAE183, A-F), JCAL171 (G-L) and CNPAE218 (M-R), subjected for 750 mM NaCl by 50 hours (stressed) and recovery after 35 days after salt stress alleviation (after Stress). ADA, Adaxial surface; ABA, Abaxial surface. Bars = 100 μm.

**Figure 3:**
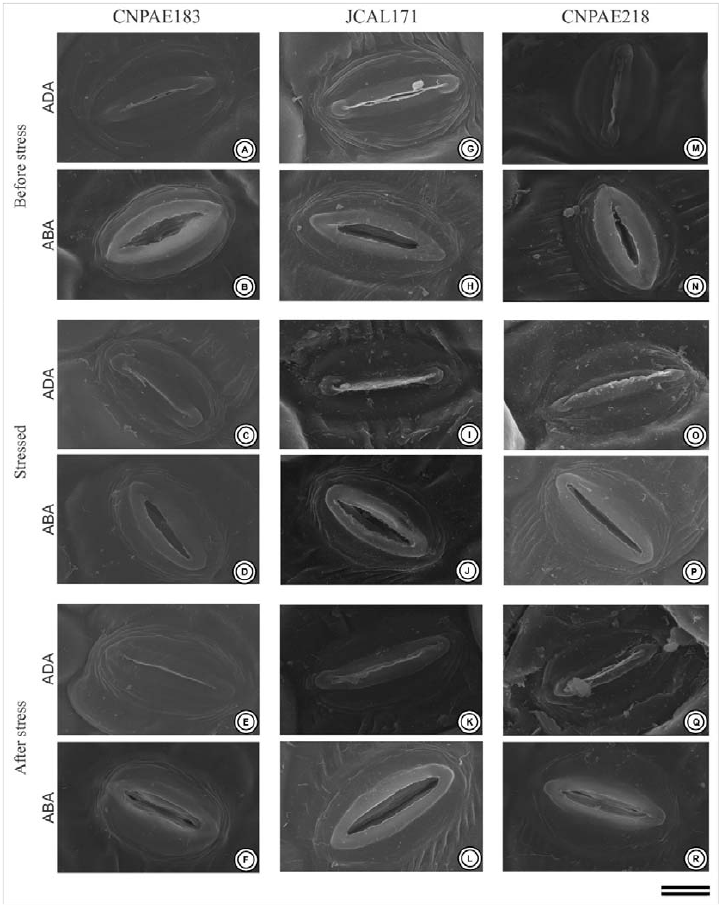
Electron scanning microscropy of the stomata of the leaves of the three different genotypes of *Jatropha curcas* (CNPAE183, A-F), JCAL171 (G-L) and CNPAE218 (M-R), subjected to 750 mM NaCl for 50 hours (stressed) and recovery after 35 days after salt stress alleviation (after Stress). ADA, Adaxial surface; ABA, Abaxial surface. Bars = 100 μm.

**Table 3.**
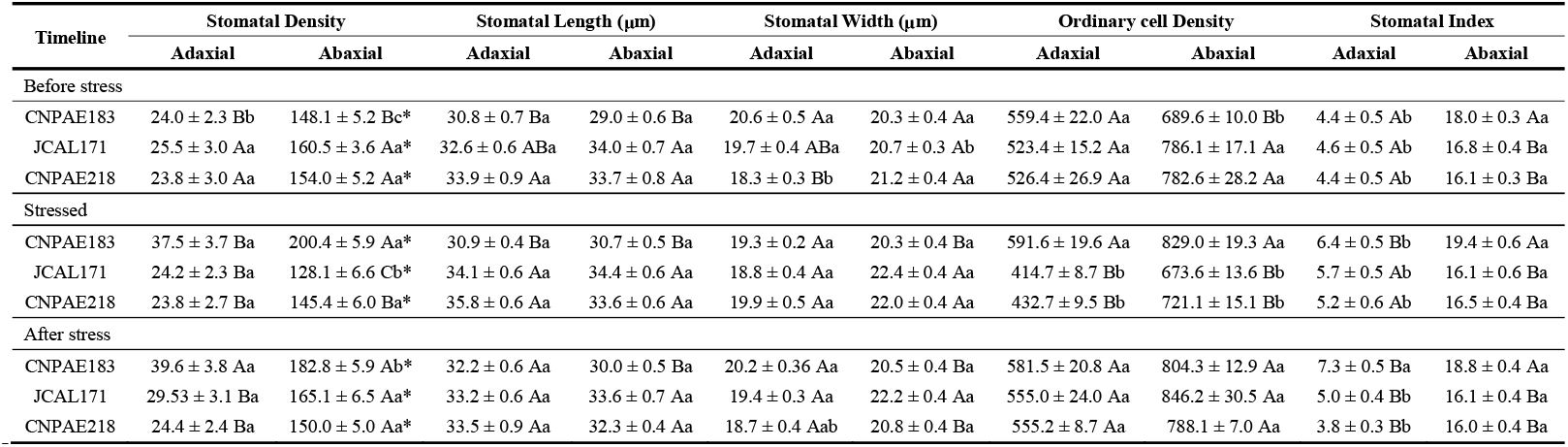
Stomatal features in the leaves of *Jatropha curcas* plants subjected to 750 mM NaCl in three events (before stress, stressed and after stress). Different upper-case letters denote statistical significance between the means for genotype and time, and different lower-case letters represent statistical significance among the genotypes in the different timeline (*P* ≤ 0.05, Newman–Keuls test). Asterisks denote statistical significance among the means in the adaxial and abaxial surface in the same stomatal feature, genotype and timeline (*P* ≤ 0.001, t-test). The values represent the media (± SE). n = 4

**Table 4.** Stomatal area and stomata aperture in leaves of *Jatropha curcas* plants subjected to 750 mM NaCl in three events (before stress, stressed and after stress). Different upper-case letters denote statistical significance between the means for genotype and time, and different lower-case letters denote statistical significance among the genotypes in the different timeline (*P* ≤ 0.05, Newman–Keuls test). Asterisks denote statistical significance among the means in the adaxial and abaxial surface in the same stomatal feature, genotype and timeline (*P* ≤ 0.001, t-test). The values represent the media (± SE). n = 4

| Timeline | Stomatal Area ($\mu\text{m}^2$) | | Stomatal Aperture ($\mu\text{m}^2$) | |
| --- | --- | --- | --- | --- |
|  | Adaxial | Abaxial | Adaxial | Abaxial |
| Before stress |  |  |  |  |
| CNPAE183 | 526.2 ± 11.3 Aa | 453.2 ± 13.7 Ba* | 12.1 ± 1.6 Aa | 31.9 ± 2.1 Ba* |
| JCAL171 | 506.3 ± 12.6 Aa | 531.1 ± 10.9 Aa | 3.3 ± 0.9 Ba | 29.9 ± 2.4 Ba* |
| CNPAE218 | 484.1 ± 11.8 Ab | 539.7 ± 19.3 Aa* | 7.5 ± 2.6 ABa | 44.2 ± 3.4 Aa* |
| Stressed |  |  |  |  |
| CNPAE183 | 480.2 ± 9.0 Ba | 469.4 ± 7.1 Ba | 3.4 ± 0.9 Ab | 25.8 ± 1.7 Ba* |
| JCAL171 | 518.2 ± 8.3 ABa | 577.3 ± 15.8 Aa* | 1.3 ± 0.5 Aa | 38.4 ± 2.6 Aa* |
| CNPAE218 | 548.0 ± 13.4 Aa | 553.7 ± 13.8 Aa | 2.9 ± 1.1 Aa | 37.2 ± 2.8 Aab* |
| After stress |  |  |  |  |
| CNPAE183 | 527.3 ± 12.9 Aa | 477.7 ± 9.9 Ba* | 9.0 ± 1.8 Aa | 30.0 ± 1.8 Aa* |
| JCAL171 | 504.2 ± 10.3 Aa | 558.9 ± 16.1 Aa* | 2.2 ± 0.7 Ba | 28.7 ± 3.2 Aa* |
| CNPAE218 | 455.7 ± 12.1 Bb | 521.4 ± 10.0Aa* | 5.6 ± 2.0 ABa | 29.3 ± 2.0 Ab* |

The ordinary cell density was significantly influenced by the salinity shock. On the adaxial surface, the CNPAE183 genotype showed no change in the density of ordinary cells under salinity, while the JCAL171 and CNPAE218 genotypes showed a respective increase of 34% and 17% in the density of the ordinary cells in the leaves that emerged after the alleviation of the saline conditions compared to those before the stress. The abaxial surface varied dramatically from adaxial surface, since the genotype CNPAE183 did not show an alteration of the density of the ordinary cells under salinity, while the genotypes JCAL171 and CNPAE218 showed a respective increase of 26% and 9.3% in the density of the ordinary cells in the leaves that emerged after the alleviation of the saline conditions in comparison to those before stress.

The stomatal index followed the same behavior as the stomatal density, where the abaxial surface had a stomatal index 3.5 times higher than that observed in the adaxial surfaces (Table 3). However, the stomatal index was the characteristically less affected by the salinity shock, since it did not demonstrated any significant difference between the means shown in the leaves before and after stress. The genotype CNPAE183 is an exception, which demonstrated a 45% increase in the stomatal index on the adaxial surface of the epidermis, but without a significant increase in the stomatal index at the abaxial surface.

## Discussion

It is noteworthy that salinity causes reduced water availability, affecting plant performance, since it interferes with the ionic balance of the cells, causing physiological, anatomic and molecular damages, permanent lesions and cell death [14, 34]. Of these features, foliar abscission and the reduction of stomatal openings are more common strategies utilized by plants to reduce their evapotranspiration [8, 9]. These findings were clearly demonstrated in this paper, where there was a strong reduction in the growth and development of the plants and a strong reduction in the stomatal opening in the order to cope salinity shock.

Melo, da Cunha [35] describes that the leaf and roots morphoanatomy of *J. curcas* were modified when submitted to high salt concentrations. According to these authors, the palisade parenchyma was demonstrated to be bi-stratified, different from what was observed in this paper where the palisadic parenchyma consisted of a single layer of compact and elongated cells. Oliveira, Pereira [36] complements this research by demonstrating that changes in the anatomy of *J. curcas*, including the xylem vessels, favored a greater or lesser drought tolerance of the genotypes when the native or potential hydraulic conductivity were analysed following water deficiency. The effects of water restriction on the root and leaf anatomy of *J. curcas* are very well studied. However, the effects of salt stress on leaf anatomy have received meager attention in this species. Therefore, we looked for examples in other species. Following salt stress, leaves of *Sporobolus ioclados* are smaller and thicker with increased epidermis thickness, and well-developed bulliform cells, and an increased mesophyll area and trichome density under high concentration of salt [37]. In *Lycium* species, low and moderate concentrations of NaCl induced variation in the leaf, mesophyll and epidermis thickness [38]. A pioneer assay in this field was conducted by Longstreth and Nobel [16]. These authors studied the effect of salinity on three plants with different responses to salt stress: *Phaseolus vulgaris*, salt-sensitive; *Gossypium hirsutum*, moderately salt-tolerant; and *Atriplex patula*, salt-tolerant. To study the changes in leaf anatomy, these authors measured different parameters, such as the epidermal, mesophyll and leaf thickness; the length and diameter of the palisade cells; the diameter of spongy cells; the surface area of the mesophylls per unit of leaf surface area; the ratio of the mesophyll cell surface area to the leaf surface area; and leaf succulence. The salt-tolerant species, *Atriplex patula*, irrigated with different NaCl solutions (0.05, 0.1, 0.2, 0.3 and 0.4 M), showed more leaf thickness due to an increase in epidermal and mesophyll thickness. Finally, major increases were observed in the succulence values. Opposite effects were observed in the other two species that were less tolerant to salt irrigation. Acosta-Motos, Ortuño [39] studied leaf anatomical changes in *Myrtus communis* and *Eugenia myrtifolia* plants subjected to a NaCl solution of 8 dS m^−1^ for the same exposure time (30 days). In *M. communis*, no significant changes were observed in the palisade parenchyma, but a decrease in the spongy parenchyma and an increase in the intercellular spaces were observed. In *Eugenia* leaves, the same anatomical changes were observed. In addition, *Eugenia* leaves displayed a remarkable increase in palisade parenchyma. These traits in *Eugenia* plants improve CO2 diffusion, making it easier for CO2 to reach the chloroplasts, which have a greater presence in the palisade parenchyma. These alterations can protect and improve the photosynthetic performance in *Eugenia* plants, especially in a situation of reduced stomatal aperture. Similar anatomical changes were also described by Rouphael, de Micco [40] in *Ecklonia maxima* grown under saline conditions. In this experiment, salt stress was accompanied by an increase in mesophyll thickness due to an increased in the intercellular spaces. Such anatomical modifications in the leaves of salt-stressed *E. maxima* plants are consistent with previous findings in the ornamental shrubs *M. communis* and *E. myrtifolia* [39].

Our results corroborated in great part of those results described above, because the leaf morphology of *J. curcas* plants affects its morphoanatomy in a similar form in response to salinity shock. The leaves of *J. curcas* subjected to salinity shock tended to be thicker than the leaves collected before stress. A tendency to increase the leaf thickness with a significant reduction of air spaces can translate into more succulent leaves that provide a smaller path for the CO_2_ from the stomatal chamber to the Rubisco carboxylation sites as previously demonstrated for many plant species [41]. Based on this, we hypothesize that *J. curcas* may reduce its air spaces in the mesophyll as a strategy to mitigate the low gas exchange promoted by the stomata closing and improve photosynthesis, increasing its water use efficiency, as previously reported by others [7, 8, 15]. Among the genotypes evaluated in this study, we highlight CNPAE183, which tended to increase the thickness of the mesophyll during the stress time, without a true translation of this thickness in the leaves that emerged after salinity alleviation but, when accompanied by the reduction of the intercellular spaces, may constitute a strategy of this genotype to favor the improve of gas exchanges and diffusion of CO2 through the mesophyll [40]. It is probable that these anatomical variations were caused by exposure to salinity shock in a relatively short time, promoting a more succulent leaves, which are associated with the accumulation of osmotically active solutes to maintain the cellular turgor pressure, mitigating the salt concentration in the mesophyll and minimizing the toxic effects of excessive ion accumulation [42].

During the entire experiment, under salinity shock, the maintenance of calcium (Ca) crystals on the leaves of all the genotypes evaluated was observed, a result that differs from the studies of Melo, da Cunha [35], which describes the disappearance of the drusen in the leaf mesophyll of *J. curcas* under salinity. The maintenance of the drusen on the leaf mesophyll as a function of salinity, described in this paper, can be explained by the low duration of exposure to the salt stress (*i.e*., salinity shock) or by the fact that in this study, the plants were irrigated with Hoagland solution plus NaCl and not only saline water as done in Melo, da Cunha [35]. Thus, the presence of Ca in the nutrient solution may have contributed to the presence of the drusen or to their maintenance in the leaf mesophyll, since Ca is immobile in the leaf tissue. Recent studies indicate that these Ca crystals, when metabolized, can supply the plant with an extra source of Ca and carbon [43, 44], essential elements to handle abiotic stress, with Ca being an important nutrient for the full development of the plant and the stabilization of cell membranes [45], while an extra carbon source could maintain photosynthesis for a time, even under conditions of high salt stress and reduced stomatal aperture [44]. It should be emphasized that saline stress led to a strong reduction in the stomatal opening of all the genotypes described in this study.

The ability of the plant to withstand abiotic stress is a complex process involving different pathways, such as the plant’s phenological stage, genetics, environmental, saline concentration and time of exposure to stress [46]. Intraspecific genetic differences have an important potential to select tolerance mechanisms of the most diverse environmental stimuli, such as saline stress and water deficit. These phenotypic variations may be related, among other things, to morphoanatomical adaptations that optimize water use efficiency mechanisms [8, 15, 46, 47], such as variations in stomatal size and density [35]. Changes in the size and distribution pattern of the stomata in *J. curcas* leaves were recently observed [47] where the authors compared *J. curcas* during the dry and rainy seasons and identified that in the dry season the leaves had a lower leaf area and higher stomatal density compared to those from the rainy season. In this paper, different genotypes of *J. curcas* with contrasting salinity tolerance were evaluated for their epidermal ultrastructure, and the genotype CNPAE183 showed stomatal cells with smaller dimensions and a higher stomatal index when compared to the genotypes JCAL171 and CNPAE218. Smaller stomata have faster dynamic characteristics [48, 49], indicating a greater control of the opening of the stomatal pore, a fact that promotes a stomatal opening/closure even with small variations in the physiological conditions. More responsive stomatal cells contribute to an optimization in the mechanism of water use efficiency [47–49] and, when associated with a higher stomatal density, may promote an increase in the photosynthetic capacity and a reduction in the effects of saline stress. In this manner, the epidermal ultrastructural features bring important information about the different mechanisms of stress tolerance, becoming a potential tool in the selection of elite genotypes with improved agricultural qualities [14, 48].

## Conclusions

Salinity tolerance is complex and involves physiological processes, but progress has been made in studying the mechanisms underlying a plant’s response to salinity. However, several previous studies on salinity tolerance could have benefitted from improved experimental design. The data from this research suggest that the salt-stressed plants modulate the expansion of stomata differently compared with other epidermal cells, while the cell division processes remain unchanged. However, changes in stomatal area have a more direct effect on the regulation of stomatal conductance and transpiration than do changes in ordinary epidermal cells. Thus, we argue that the genotype CNPAE183 demonstrates indications that it can be considered to be as a promising genotype for future studies of genetic improvement that involve the search for elite genotypes tolerant to salinity. We hope that this paper will provide pertinent information to researchers on performing proficient assays and interpreting the results from salinity tolerance experiments.

## Supplementary data

Plant architecture of CNPAE183 (A and B), JCAL171 (C and D) e CNPAE218 (E and F) of *Jatropha curcas* genotypes subjected to the control (A, C e E) and salt stressed (B, D e F). Down panel: morphotype of leaves of CNPAE183 (G), JCAL171 (H) and CNPAE218 (I), de *Jatropha curcas* morphotypes. LA, leaf area; SLA, specific leaf area. Upper and lower bars 10 cm and 5 cm, respectively. The values represent the media (± SE). n = 20

## Acknowledgments

The authors thanks the National Council for Scientific and Technological Development (CNPq Grants 404357/2013-0). Leonardo Silva-Santos and Natália Corte-Real thanks the Foundation for Science and Technology of Pernambuco, FACEPE (Grants IBPG-1799-2.07/2015 and IBPG-0709-2.07/13, respectively) to scholarship. The authors would also like to thank Dr. Karina Lidianne Alcântara Saraiva (Technology Platforms Nucleus, Aggeu Magalhães Institute) and MsC. Ana Carla da Silva (Cell Biology of Pathogens Laboratory, Aggeu Magalhães Institute) for they help with scanning electron microscopy procedures. The authors would also like to anonymous reviewers for their kind revisions of this manuscript.

---

**Supplementary Figure 1:**
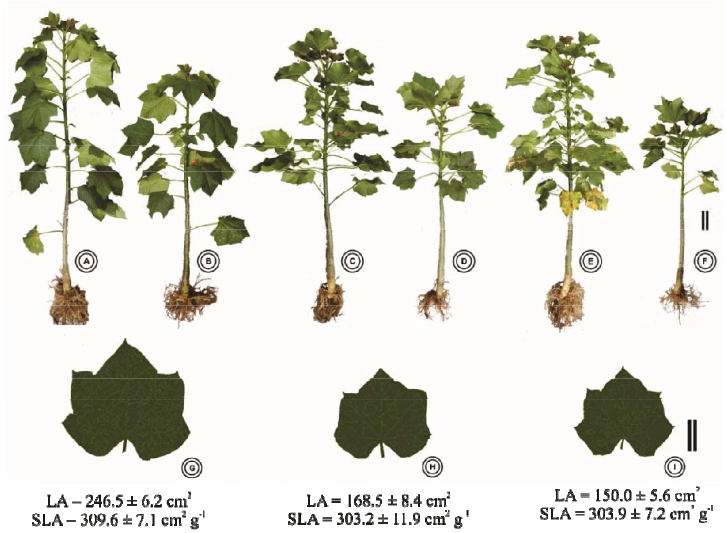
Upper panel: Plant architecture of CNPAE183 (A and B), JCAL171 (C and D) e CNPAE218 (E and F) of *Jatropha curcas* genotypes subjected to the control (A, C e E) and salt stressed (B, D e F). Down panel: morphotype of leaves of CNPAE183 (G), JCAL171 (H) and CNPAE218 (I), de *Jatropha curcas* morphotypes. LA, leaf area; SLA, specific leaf area. Upper and lower bars 10 cm and 5 cm, respectively. The values represent the media (± SE). n = 20

